# IL-27 skews TNF-alpha-induced inflammatory microenvironment in keratinocytes

**DOI:** 10.1101/2022.03.18.484839

**Authors:** Akihiro Aioi, Tomozumi Imamichi

## Abstract

Inflammaging has received considerable attention because aging is characterized by low-grade, chronic and asymptomatic inflammation, concomitant with increased blood levels of senescence-associated secretory phenotype (SASP) factors, including IL-1, IL-6, IL-8, IL-18 and tumor necrosis factor-alpha (TNF-alpha). On the other hand, IL-27 is not categorized as SASP factors though it is known that IL-27 has pleiotropic roles in inflammation. Here, we evaluated the interaction between TNF-alpha and IL-27 in the context of low-grade inflammation by using in HaCaT cells. TNF-alpha induced significant upregulation of IL-6 and IL-8 through the experimental concentrations (~10 ng/ml) of TNF-alpha, while the mRNA expression levels of IL-1RA, IL-10 and IL-18BP were unchanged. After confirming the expression of IL-27 receptor in HaCaT cells, we examined the effects of IL-27 alone on the cytokine expression. IL-27 alone significantly enhanced mRNA expression levels of IL-10 and IL-18BP by 1.61-fold and 1.46-fold, respectively, and also enhanced mRNA expression levels of IL-6 by 2.32-fold. In the presence of 100 ng/ml IL-27, the expression levels of anti-inflammatory cytokines, IL-1RA, IL-10 and IL-18BP, were significantly upregulated with the treatment of a physiological concentration (1 ng/ml) TNF-alpha. Taken together, a high concentration of IL-27 exhibits anti-inflammatory effects in the presence of a low concentration of TNF-alpha when keratinocytes are the recipient of IL-27 signaling, suggesting the anti-inflammatory roles of IL-27 in inflammaging may be regulated by TNF-alpha concentration.

## Introduction

Interleukin (IL)-27, a member of the IL-12 family is a heterodimeric cytokine composed of IL-27 p28 and Epstein–Barr virus-induced gene protein 3 subunits. The functional IL-27 receptor consists of IL-27R alpha (WSX-1) and glycoprotein 130 (gp130), a commonly used cytokine receptor for the IL-6 family [1]. Once binding to IL-27 receptor, IL-27 activates JAK 1, 2, TYK2, STAT-1, 2, 3, 4 and 5 [2–5], followed by diverse regulatory functions of IL-27. As previous studies reported that IL-27 has pro-inflammatory and anti-inflammatory effects [6–9] and anti-viral effects [10], IL-27 displays a dual role in inflammation. In the skin, the potential roles of IL-27 have been previously reported. Yang *et al.* demonstrated that IL-27 produced by CD301b^+^ cells which infiltrated into wound lesion, stimulated keratinocyte proliferation and re-epithelialization in wound-healing process [11]. Another study reported that IL-27 was expressed in chronic lesional allergic eczematous [12]. In addition, a previous study showed that serum IL-27 levels in psoriatic patients were significantly higher than those in healthy controls, correlating with disease severity [13]. Aging becomes a serious issue in an aging society. Previous studies have reported that aging is strongly associated with inflammation [14–16], which is the basis of the concept of inflammaging [17]. Clinically, inflammaging consists of low-grade, chronic, and asymptomatic inflammation, characterized by increased blood levels of several inflammatory biomarkers, including IL-1, IL-6, IL-8, IL-18, and tumor necrosis factor-alpha (TNF-alpha), which are together called senescence-associated secretory phenotype (SASP) factors [18–20]. The SASP factors secreted from senescent cells, which are considered passive bystanders, play pivotal roles in aging. Skin aging can be divided into two distinct types, intrinsic aging and extrinsic aging, based on the fact that the skin is the outermost organ and is therefore exposed to the external environment. However, despite their different histological features and functional triggers, intrinsic and extrinsic aging share common biochemical mechanisms, involving SASP factors [21]. Here, we explored the effect of IL-27 on the inflammatory environment of HaCaT cells, a human-derived keratinocyte cell line.

## Materials and Methods

### Cytokine

IL-27 and TNF-alpha were purchased from Proteintech (Rosemont, IL, USA). Both cytokines were dissolved in sterilized PBS at the concentration of 100 μg/ml and subjected to experiments.

### Cell culture

To maintain the distinct differentiation stage of HaCaT cells, calcium in fetal bovine serum (FBS) was depleted by incubation with Chelex 100 resin (BioRad, Hercules, CA, USA) for 1 h at 4 °C. HaCaT cells were maintained in Ca^2+^-free Dulbecco’s modified Eagle’s medium (DMEM) supplemented with 5% Ca^2+^-depleted FBS, 4 mM glutamine, 1 mM sodium pyruvate and 2 mM CaCl_2_ at 37 °C in a 5% CO2-humidified atmosphere.

### Immunoblotting

HaCaT cells were treated with 3, 10, 30 and 100 ng/ml of IL-27 for 20 min at 37 °C, and then subjected to immunoblotting. After washing with PBS, HaCaT cells were collected and lysed in the RIPA buffer supplemented with protease inhibitors and phosphatase inhibitors. Equal amount of protein (10 μg) were loaded, resolved via SDS-PAGE and transferred to PVDF membrane, followed by immunoblotting with phospho-STAT3 rabbit monoclonal antibody (Cell Signaling Technology, Danvers, MA, USA). Immunoreactive proteins were visualized using an enhanced chemiluminescence detection system (Millipore, Bedford, MA, USA).

### Treatment with IL-27 or (and) TNF-alpha

To estimate the effects of IL-27 on the mRNA expression levels of cytokines, HaCaT cells were treated with 10, 30 and 100 ng/ml of IL-27 for 24 h at 37 °C. To estimate the effects of TNF-α on the mRNA expression levels of cytokines, HaCaT cells were treated with 1, 5 and 10 ng/ml of TNF-alpha for 24 h at 37 °C. To estimate the interaction of TNF-alpha and IL-27 on the mRNA expression levels of cytokines, HaCaT cells were treated with 1 or 10 ng/ml of TNF-alpha for 24 h at 37 °C in the presence of 10 or 100 ng/ml of IL-27.

### Real-time qPCR

After the treatments, cells were harvested and total RNA was prepared with SV RNA isolation kit (Promega, Madison, WI, USA), according to the manufacturer’s instructions, followed by reverse transcription using ReverTra Ace® qPCR RT Master Mix (TOYOBO, Osaka, Japan). PCR amplification and detection were conducted on a CFX96 real-time PCR system (BioRad, Hercules, CA, USA) using the initial denaturation condition of 95 °C for 3 min, followed by 40 cycles at 95°C for 5 sec and 60 °C for 30 sec each. The following primer pairs were used: beta-actin, 5’-GATGAGATTGGCATGGCTTT-3’ (sense) and 5’-CACCTTCACCGTTCCAGTTT-3’ (antisense); IL-27R alpha, 5’-AGGCCACCTCACCCACTACA-3’ (sense) and 5’-ATGCTGTCACCCACAGCTCA-3’ (antisense); gp130, 5’-ATCCTGTGGATCTGGGCAAA-3’ (sense) and 5’-GCCTCCATGCCAACTGTTTC-3’ (antisense); IL-1RA, 5’-TGGAGGGAAGATGTGCCTGT-3’ (sense) and 5’-GCGCTTGTCCTGCTTTCTGT-3’ (antisense); IL-6, 5’-ATGCAATAACCACCCCTGAC-3’ (sense) and 5’-AAAGCTGCGCAGAATGAGAT-3’ (antisense); IL-8, 5’-ACCACCGGAAGGAACCATCT-3’ (sense) and 5’-TTGGCAAAACTGCACCTTCA-3’ (antisense); IL-10, 5’-CAAGCCTGACCACGCTTTCT-3’ (sense) and 5’-AAAGGGGCTCCCTGGTTTCT-3’ (antisense); IL-18BP, 5’-TGGGTTCACACGCAGCTAGA-3’ (sense) and 5’-GGCATCTGCTTCCTCCTTCA-3’ (antisense). The expression of target mRNA was quantified using the comparative threshold cycle (Ct) method for relative quantification (2^-ΔΔ^Ct), normalized to the geometric mean of the reference gene beta-actin.

### Statistical analysis

Data are expressed as means±SEM. Statistical comparison between experimental groups and controls was performed using an unpaired Student’s *t*-tests. P values less than 0.05 were considered significant.

## Results

### Functional IL-27 receptor expressed in HaCaT cells

First, real-time qPCR was conducted to estimate the expression levels of IL-27 R alpha and gp130. The expression level of IL-27R alpha mRNA was 0.21±0.05-fold of involucrin which is a differentiation maker of HaCaT cells, while the expression level of gp130 was 9.24±1.06-fold of involucrin (Fig. 1a). Next, immunoblotting for p-STAT3 was performed to confirm the function of IL-27 receptor expressed in HaCaT cells. The spot of p-STAT3 was detected in HaCaT cells treated with IL-27 through the concentration of 3 to 100 ng/ml (Fig. 1b).

**Figure 1.**
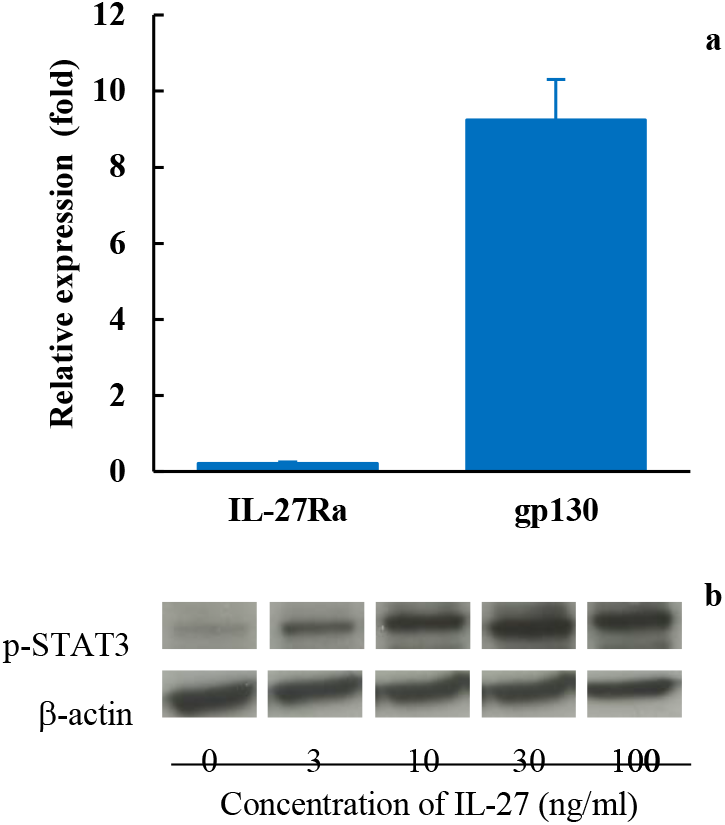
The functional IL-27 receptor is expressed in HaCaT cells. To evaluate the functional IL-27 receptor expression in HaCaT cells, qPCR (a), and immunoblotting of p-STAT3 (b) were performed. Significant mRNA levels of IL-27Rα and gp130 against IVL were confirmed in HaCaT cells (a). The spots of p-STAT3 in the immunoblot of cells treated with IL-27 were detected within a concentration range of 3 to 100 ng/ml of IL-27 (b).

### The treatment of TNF-alpha induced the mRNA expression levels of proinflammatory cytokines in HaCaT cells

To evaluate the effects of TNF-alpha on the mRNA expression levels of cytokine, the mRNA expression levels of IL-6, IL-8, IL-1RA, IL-10 and IL-18BP examined by real-time qPCR. The expression levels of IL-6 were significantly enhanced to 1.56±0.21-fold and 1.37±0.12-fold with the treatment of 5 and 10 ng/ml TNF-alpha respectively. The expression of levels of IL-8 were significantly upregulated up to 8.43±1.07-fold with the treatment of 10 ng/ml TNF-alpha. The expression levels of IL-1RA, IL-10 and IL-18BP were not affected with the treatment of TNF-alpha (Fig. 2).

**Figure 2.**
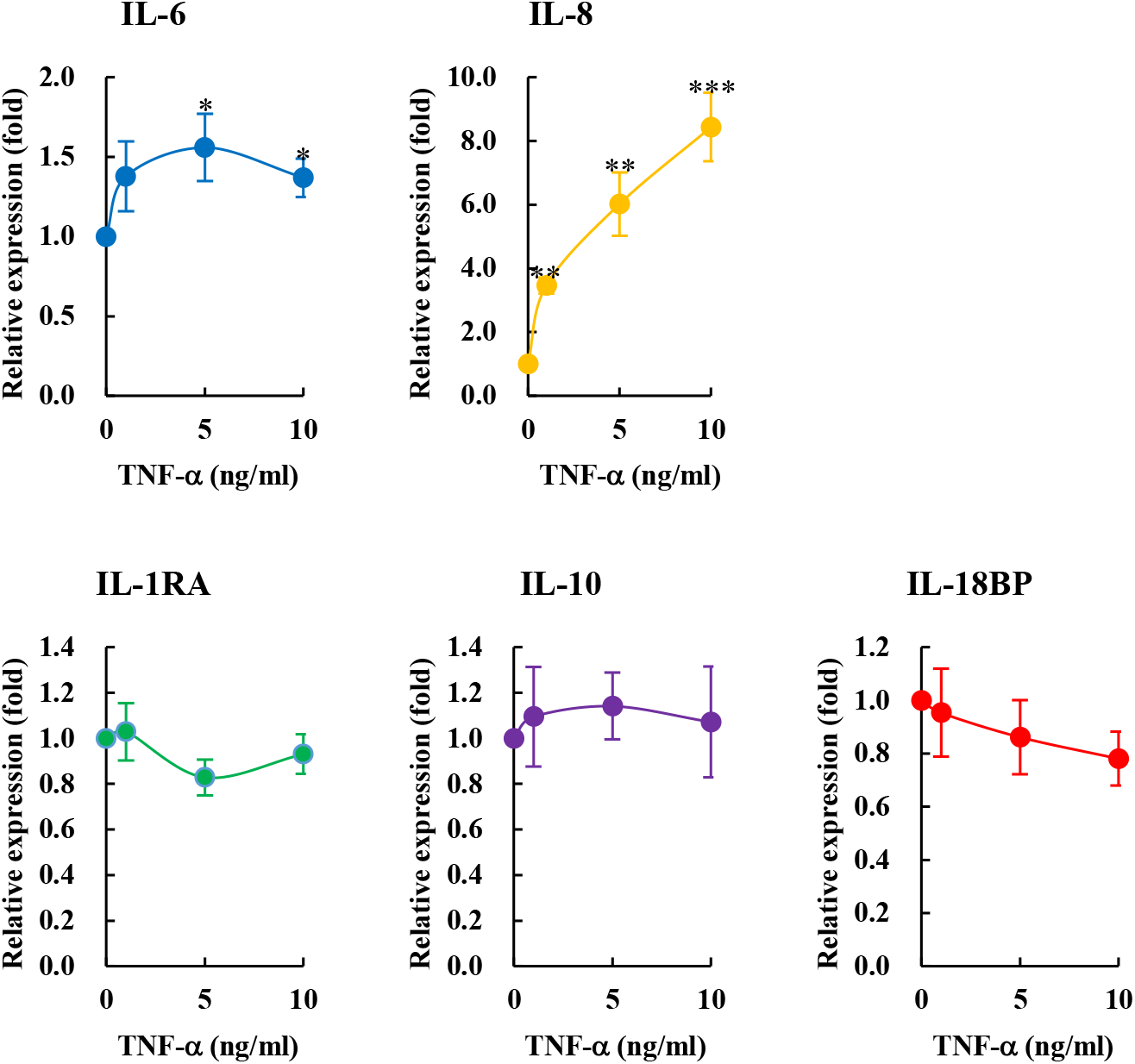
TNF-α-induced upregulation of IL-6 and IL-8 mRNA expression. To evaluate the effect of TNF-α, the mRNA levels of IL-6, IL-8, IL-1RA, IL-10, and IL-18BP were measured in HaCaT cells. TNF-α stimulation upregulated the mRNA levels of IL-6 and IL-8, while the those of IL-1RA, IL-10, and IL-18BP remained unchanged at experimental concentrations. Each value is represented as the mean±SEM of at least three independent experiments (*p<0.05, ***p<0.001 compared with the control).

### The treatment with IL-27 affected the mRNA expression levels of IL-6, IL-10 and IL-18BP

To evaluate the effects of IL-27 on the mRNA expression levels of cytokine, the mRNA expression levels of IL-6, IL-8, IL-1RA, IL-10 and IL-18BP examined by real-time qPCR. The expression levels of IL-6 were significantly enhanced up to 2.32±0.10-fold with the treatment of 100 ng/ml IL-27, with dose-relation manner. The expression level of IL-10 was significantly upregulated to 1.61±0.23-fold with the treatment with 100 ng/ml IL-27. The expression level of IL-18BP was significantly upregulated with the treatment with 10 ng/ml IL-27. The expression levels of IL-8 and IL-1RA were not affected with the treatment of IL-27 (Fig. 3).

**Figure 3.**
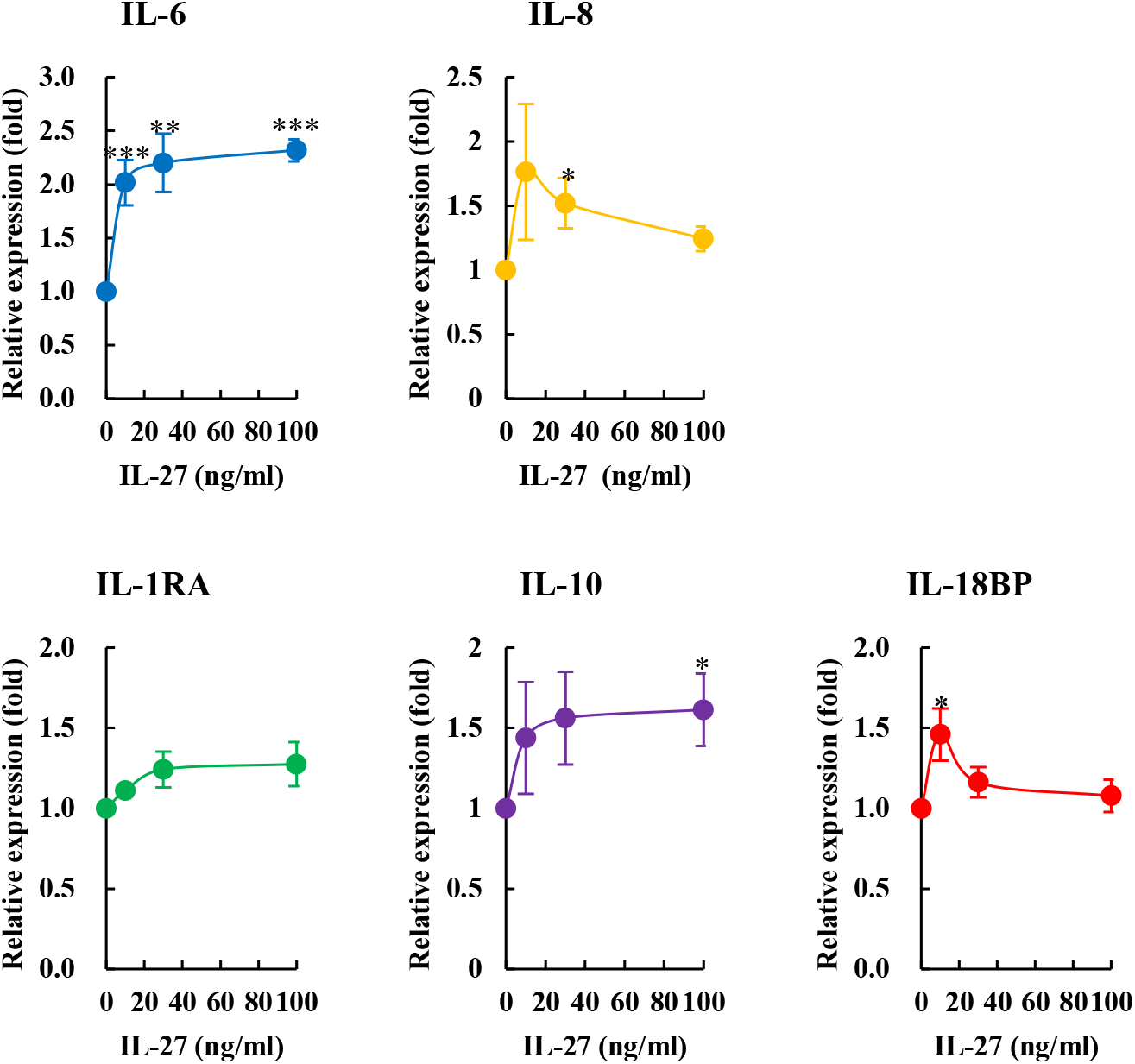
IL-27 induced the upregulation of mRNA expression of IL-6, IL-10 and IL-18BP. To evaluate the effect of IL-27, the mRNA levels of IL-6, IL-8, IL-1RA, IL-10 and IL-18BP were measured in HaCaT cells. The mRNA expression of IL-6 was significantly upregulated in a dose-dependent manner. The mRNA expression of IL-10 and IL-18BP was also upregulated at concentrations of 30 ng/ml and 10ng/ml respectively. Each value is represented as the mean±SEM from at least three independent experiments (*p<0.05, **p<0.01, ***p<0.001 compared with the production from the control).

### TNF-alpha affected IL-27-induced expression levels of cytokines

To estimate the interaction of TNF-α and IL-27 on the mRNA expression levels of cytokines, IL-27-regulated expression levels were determined in the presence of 1 or 10 ng/ml TNF-α. In the presence of a low dose IL-27 (10 ng/ml), the expression levels of IL-10 and IL-18BP were significantly upregulated with the treatment of 10 ng/ml TNF-α, while the expression levels of IL-6 and IL-8 were significantly enhanced (Fig. 4). On the other hand, in the presence of a high dose IL-27 (100 ng/ml), the expression levels of anti-inflammatory cytokines, IL-1RA, IL-10 and IL-18BP, were significantly upregulated to 1.70±0.0.16-fold, 3.30±0.28-fold and 2.87±0.49-fold, resupectively, with the treatment of 1 ng/ml TNF-α, while the expression levels of proinflammatory cytokines also upregulated (Fig. 5).

**Figure 4.**
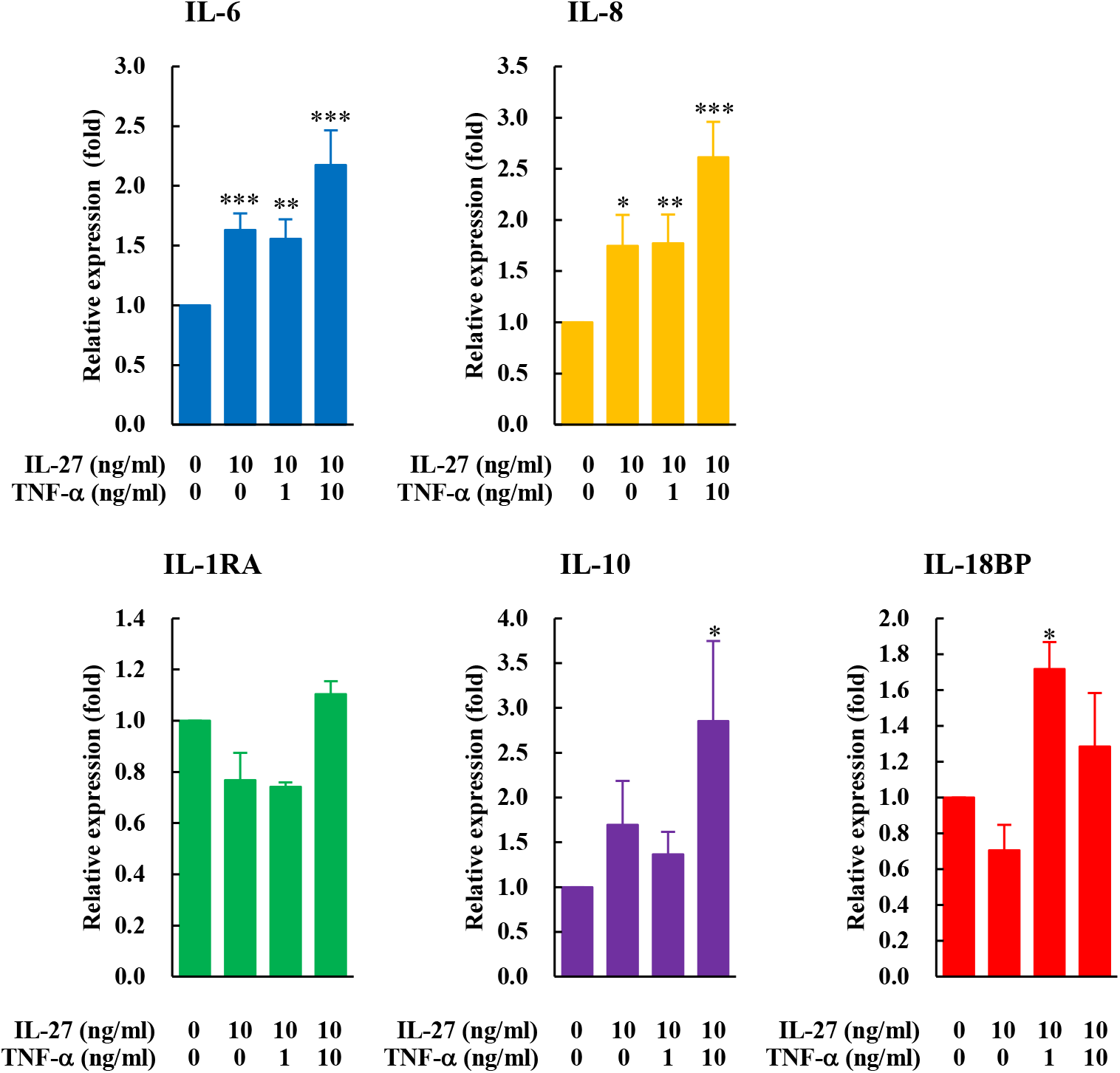
The interaction between IL-27 and TNF-α in the presence of 10 ng/ml IL-27. To examine the effect of interaction between TNF-α and IL-27 on the mRNA levels of cytokines, IL-27-regulated mRNA levels were determined in the presence of 1 or 10 ng/ml TNF-α. In the presence of 10 ng/ml IL-27, the mRNA levels of IL1RA and IL-18BP were significantly upregulated by TNF-α, while the those of other cytokines were only slightly upregulated. Each value is represented as the mean± SEM from at least three independent experiments (*p<0.05, **p<0.01, ***p<0.001 compared with the production from the control).

**Figure 5.**
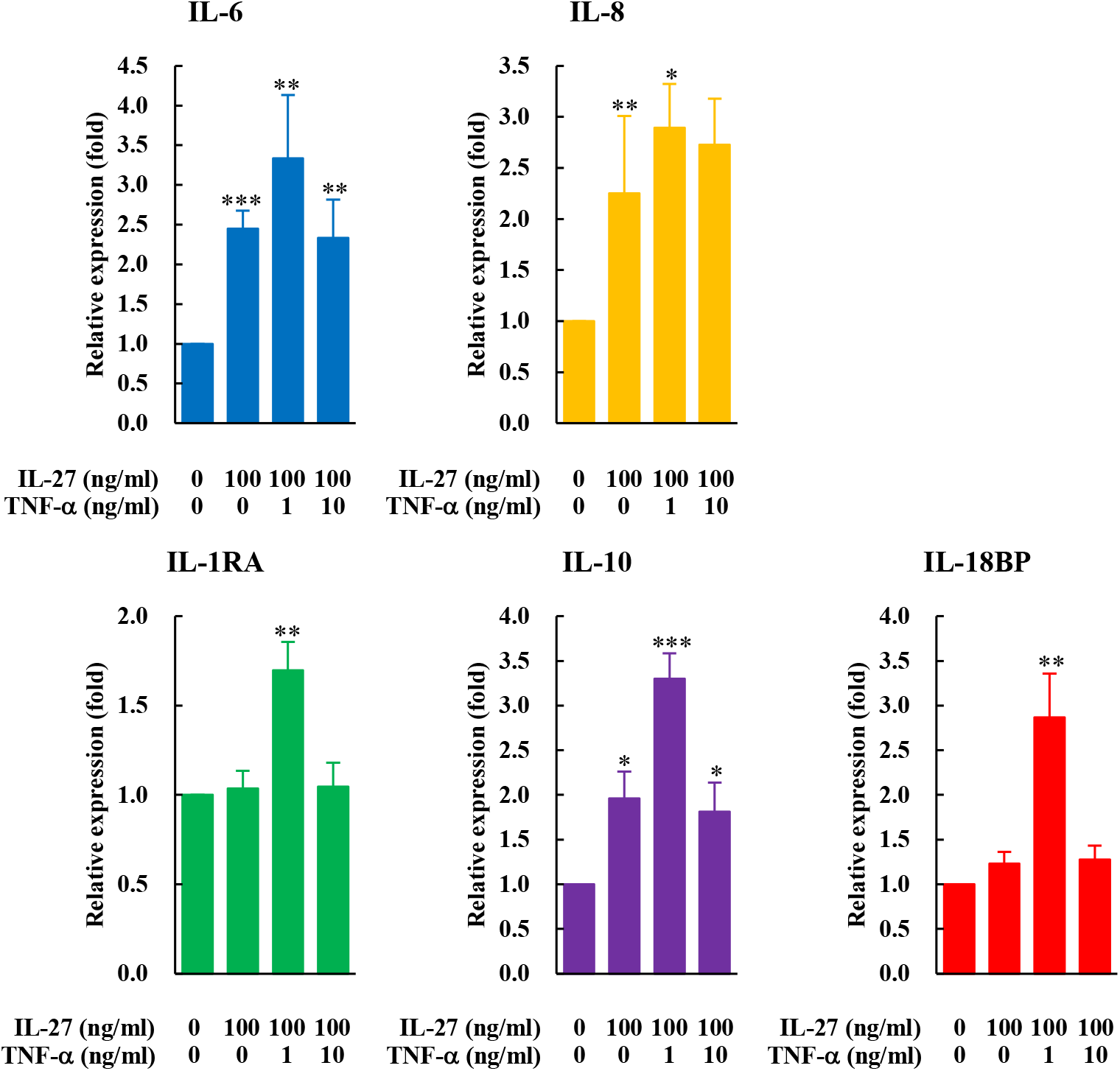
The interaction between IL-27 and TNF-α in the presence of 100 ng/ml IL-27. To examine the effect of interaction between TNF-α and IL-27 on the mRNA levels of cytokines, IL-27-regulated mRNA levels were determined in the presence of 1 or 10 ng/ml TNF-α. In the presence of 100 ng/ml IL-27, the mRNA levels of IL1RA, IL-10 and IL-18BP were significantly upregulated in response to treatment with 1ng/ml TNF-α, while those of pro-inflammatory cytokines were not changed. Each value is represented as the mean±SEM from at least three independent experiments (*p<0.05, **p<0.01, ***p<0.001 compared with the production from the control).

## Discussion

The aging of organs starts at the time of birth and continues throughout life. How to cope with aging, which causes systemic as well as topical frailty, is a growing subject of interest because of the aging society. Previous studies have reported strong association between inflammation and aging [14–16], which is the basis of the concept of inflammaging [17]. In inflammaging, which is clinically characterized by low-grade, chronic, and asymptomatic inflammation [18, 19], SASP factors such as IL-1, IL-6, IL-8, and TNF-α play a pivotal role in the progression of aging. Among these SASP factors, TNF-α, an immunological biomarker of frailty and aging [15, 22], is known to play a vital role in inflammatory diseases such as psoriasis [22]. In addition, a previous study has reported that TNF-α induces the production of IL-27 from human macrophages [23]. Moreover, other previous studies have demonstrated that IL-27, whose serum levels in patients with psoriasis were elevated, suppressed the TNF-α-induced production of IL-1α and CCL20 [13]. Moreover, as per another study, IL-27 suppressed the TNF-α-induced production of CXCL1, CXCL2, CXCL8, and CCL20 from human keratinocytes [24]. These previous studies suggest that the interaction between TNF-α and IL-27 plays a vital role in psoriasis. Here, using HaCaT cells, a human-derived keratinocyte cell line, we explored how IL-27 affects the TNF-α-induced inflammatory environment although IL-27 is not categorized as an SASP factor. We evaluated the expression and function of IL-27 receptor in HaCaT cells, as to the best of our knowledge, it has not been evaluated before. The results of qPCR and immunoblotting demonstrated that both subunits of the IL-27 receptor, IL-27Rα and gp130, were expressed in HaCaT cells, and that p-STAT3 was present in IL-27-treated HaCaT cells. These results suggest that a functional IL-27 receptor is expressed in HaCaT cells. We also examined the effects of TNF-α on cytokine production in HaCaT cells. As shown in Fig. 2, the treatment with TNF-α upregulated the mRNA levels of IL-6 and IL-8, while those of IL-1RA, IL-10 and IL-18BP were not affected. In agreement with previous studies, this study demonstrated the pro-inflammatory effects of TNF-α. Next, we evaluated the effects of IL-27 alone on the mRNA levels of IL-6, IL-8, IL-1RA, IL-10, and IL-18BP (Fig. 3). Surprisingly, IL-27 upregulated IL-6 mRNA expression in a dose-dependent manner. Although Kalliolias *et al.* demonstrated that the priming of human macrophages with IL-27 enhanced IL-6 production by stimulation with toll-like receptor ligands, while IL-8 production induced by TNF-α was suppressed in the presence of IL-27 [25], to the best of our knowledge, the present study is the first to demonstrate that IL-27 alone can induce IL-6 production. Regarding the effects of IL-27 as an anti-inflammatory cytokine, IL-27 significantly upregulated the mRNA expression of IL-10 and IL-18BP at concentrations of 30 ng/ml and 10 ng/ml, respectively, which is in agreement with the findings of previous studies [26, 27]. These results suggest that IL-27 exerts both pro- and anti-inflammatory effects on keratinocytes. Next, we evaluated the effects of IL-27 and TNF-α on cytokine production. To the best of our knowledge, although 10 ng/ml of TNF-α has been used in vitro to mimic symptomatic skin inflammation of the type observed in psoriasis, there has been no previous study that used 1ng/ml of TNF-α. We speculate that the reason why a low concentration of TNF-α has not been used so far is that it does not produce clear results with respect to cytokine production. However, because the elevated plasma concentration of TNF-α (below 10 pg/ml) in the elderly is associated with frailty and mortality, it is likely that low-dose TNF-α plays a role in inflammaging [15, 28]. Thus, we used concentrations of 1 and 10 ng/ml of TNF-α in this study. We observed different expression patterns of cytokines depending on the concentration of IL-27. In the presence of high doses of IL-27 (100 ng/ml), the mRNA levels of anti-inflammatory cytokines were significantly upregulated when the cells were treated with 1 ng/ml TNF-α (Fig.5), whereas the expression levels of IL-1RA and IL-10 were significantly upregulated with the treatment of 10ng/ml TNF-α in the presence of low dose of IL-27 (10ng/ml), as shown in Fig. 4. Collectively, the findings of this study indicate that at high concentrations, IL-27 exhibits anti-inflammatory effects in the presence of low concentrations of TNF-α in keratinocytes, indicating its anti-inflammatory role in inflammaging. However, because it has been reported in a previous study that TNF-α signaling via IL-27R promotes the aging of hematopoietic stem cells [29], and it remains unclear whether sufficient amount of IL-27 is supplied by dendritic cells and macrophages, which are major sources of IL-27 whose functions decline with aging [30, 31], further experiments are required.

## Acknowledgements

The authors thank Dr. Ryuta Muromoto and Dr. Jun-ichi Kashiwakura for their valuable suggestions regarding this study. We would like to thank Editage (www.editage.com) for the English language editing. Finally, the authors appreciate Professor Tadashi Matsuda for helping with this study.

